# HSFA1 proteins mediate heat-induced accumulation of CPT7-derived polyprenols affecting thylakoid organization

**DOI:** 10.1101/2021.12.22.473876

**Authors:** Daniel Buszewicz, Łucja Kowalewska, Radosław Mazur, Marta Zajbt-Łuczniewska, Liliana Surmacz, Katarzyna Sosnowska, Renata Welc, Małgorzata Jemioła-Rzemińska, Paweł Link-Lenczowski, Agnieszka Onysk, Karolina Skorupinska-Tudek, Hsiang-Chin Liu, Yee-Yung Charng, Rafał Archacki, Wieslaw I. Gruszecki, Ewa Swiezewska

## Abstract

Polyprenols are ubiquitous isoprenoid compounds that accumulate in large quantities in plant photosynthetic tissues. While our knowledge of polyprenol biochemistry is constantly expanding, the regulation of their biosynthesis as well as the molecular basis of their cellular action are still poorly understood. In Arabidopsis, the polyprenols Pren-9, −10 and −11, synthesized by *cis*-prenyltransferase 7 (CPT7), are localized in plastidial membranes and affect the photosynthetic performance of chloroplasts.

In this report we present evidence that plastidial polyprenols are among the major constituents of thylakoid membranes. Disturbances in polyprenol level, caused by alterations in *CPT7* expression, change chloroplast ultrastructure, affect aggregation of LHCII complexes and modulate non-photochemical quenching (NPQ). Moreover, we show that Arabidopsis responds to high temperature by upregulating expression of *CPT7* and increasing the accumulation of CPT7-derived polyprenols. These heat-induced changes in polyprenol biosynthesis are mediated by Heat Shock Transcription Factors of the HSFA1 family, the master regulators of heat stress response. Collectively, results presented in this report bring us closer to understanding the mechanisms by which polyprenols affect plant physiology and provide an additional link between chloroplast biology and plant responses to changing environmental conditions.

**One sentence summary:** Heat Shock Transcription Factors induce biosynthesis of polyprenols - isoprenoid compounds that affect the organization and function of chloroplasts.

## Introduction

*cis*-Prenyltransferases (CPTs) are ubiquitous enzymes present in all living organisms. They are responsible for the biosynthesis of polyprenols – lipid molecules composed of linearly assembled isoprene residues (Swiezewska and Danikiewicz, 2005). CPTs sequentially add isopentenyl diphosphate (IPP) molecules to an allylic prenyl diphosphate starter molecule, generating double carbon-carbon bonds in *cis* geometry. The resulting polyisoprenoid chain can range from several isoprene units to thousands in natural rubber. Some plant polyprenols of a species-specific profile tend to accumulate in large quantities in leaves (Surmacz and Swiezewska, 2011), while others are converted into dolichols – essential metabolites involved in protein glycosylation – by saturation of the OH-terminal isoprene residue (Cantagrel et al, 2010; Jozwiak et al, 2015). The common feature of all isoprenoids, including polyprenols, is their biosynthetic origin from two compounds: isopentenyl diphosphate (IPP) and its isomer dimethylallyl diphosphate (DMAPP), both of which can be obtained in plants from two sources: from the cytoplasmic mevalonic acid (MVA) pathway and from the plastidial 2-*C*-methyl-D-erythritol 4-phosphate (MEP) pathway (Rodríguez-Concepción and Boronat, 2002). The compartmentalization of IPP / DMAPP biosynthesis leads to two groups of end products: cytoplasmic MVA-derived compounds (e.g. sterols) and MEP-derived metabolites of plastidial origin (e.g. phytol, carotenoids, gibberellins, abscisic acid). However, some limited exchange of IPP between cytoplasm and plastids has been reported, resulting in the synthesis of isoprenoids of mixed origin (summarized in Lipko and Swiezewska, 2016).

In plants, CPTs are encoded by gene families (Surmacz and Swiezewska, 2011; Akhtar et al, 2013) with the Arabidopsis genome containing nine putative *CPT* genes (*CPT1-9*), of which only *CPT1*, *CPT6* and *CPT7* have been partially characterized. *CPT1* is expressed exclusively in roots and the CPT1 enzyme localizes in the ER and was shown to synthesize a family of long-chain polyprenols, ranging from Pren-19 to Pren-24, that are further reduced to dolichols and affect the development of the root system (Cunillera et al, 2000; Oh et al, 2000; Surowiecki et al, 2019). On the other hand, CPT6 is also a root-specific enzyme and is involved in the biosynthesis of a single, short polyprenol, Pren-7, of unknown biological function (Kera et al, 2012; Surmacz et al, 2014). CPT7 is a plastidial enzyme utilizing geranylgeranyl diphosphate (GGPP) and IPP to synthesize polyprenols (Pren-9, −10 and −11) that are dominant polyprenol species in Arabidopsis and reside mainly in thylakoid membranes (Akhtar et al, 2017). In contrast to the products of CPT1 which are used predominantly as substrates for dolichol biosynthesis, the CPT7-derived polyprenols remain unsaturated and accumulate in relatively large quantities in photosynthetic tissues. Recently, it has been shown that plastidial polyprenols affect photosynthetic performance as plants deficient in polyprenol synthesis had decreased steady-state quantum yield of photosystem II (Y(II)) (Akhtar et al, 2017). These plants also displayed a reduced rate of electron transport and decreased photochemical quenching parameter (qL), showing that polyprenol deficiency negatively affects electron transfer from photosystem II (PSII) to the cytochrome *b_6_f* complex. The results of fluorescence anisotropy measurements suggested that polyprenols promote the formation of a more rigid structure of thylakoid membranes. Therefore, polyprenols might act as regulators of thylakoid membrane dynamics, affecting the diffusion of electron carriers and possibly affecting the organization of chloroplast complexes (Akhtar et al, 2017; Van Gelder et al, 2018). Interestingly, the effects exerted by polyprenols *in planta* were contradictory to those noted *in vitro* for model phospholipid membranes where polyprenols were observed to increase the fluidity and permeability of lipid bilayers (Hartley and Imperiali, 2012, and references therein). Recently, it was also shown that polyprenols might affect the organization of prolamellar bodies in developing Arabidopsis seedlings (Bykowski et al, 2020).

Although our knowledge of the enzymatic activity of CPTs and the biological roles of polyisoprenoid alcohols has significantly increased, mechanisms regulating the expression of *CPTs* are still poorly characterized. At the same time there are several reports showing that expression of *CPTs* and polyisoprenoid accumulation is regulated in response to environmental stimuli (Jozwiak et al, 2013; Jozwiak et al, 2017; Milewska-Hendel et al, 2017). Heat stress is one of the major environmental hazards affecting all living organisms, especially in the context of the ongoing climate changes. Adaptation of plants to high temperature is a complex, multi-level process requiring the concerted action of signaling molecules, transcriptome reprogramming, protein stabilization by chaperones, reactive oxygen species (ROS) detoxification and synthesis of protective compounds. Heat shock transcription factors (HSFs) of the HSFA1 group are master regulators of heat stress response, governing a vast network of transcriptional regulators modulating expression of a multitude of heat-responsive genes. The Arabidopsis genome encodes four members of the HSFA1 group (HSFA1A, B, D and E) (Liu et al, 2011). All four of them act redundantly as regulators of plant development and either HSFA1A, B or D alone can trigger the adaptation to heat stress (Liu et al, 2011; Yoshida et al, 2011). HSFA1s were shown to regulate the expression of multiple genes encoding heat shock proteins (HSPs), detoxification enzymes and many other proteins required for adaptation and survival in high temperature conditions (Yoshida et al, 2011; Liu and Charng, 2013). There are multiple reports showing that high temperature negatively affects photosynthesis (Hu et al, 2020). Additionally, high temperature has a profound effect on lipid membranes, increasing their fluidity and permeability (Niu and Xiang, 2018) as well as affecting their composition (Higashi and Saito, 2019; Shiva et al, 2020). Membrane lipids are essential for temperature sensing and heat stress response (Saidi et al, 2010) and, in animals, the interaction of heat shock proteins (HSPs) with specific plasma membrane lipids was suggested to be crucial for HSPs proper function (Escriba et al, 2008).

In this paper we show that *CPT7* and its products are implicated in the Arabidopsis response to heat stress since *CPT7* expression and polyprenol accumulation are induced by high temperature. Moreover, we provide evidence that heat shock transcription factors of the HSFA1 family are required for the induction of *CPT7* expression and warrant heat-induced accumulation of polyprenols. Finally, we show that polyprenols affect the ultrastructure of thylakoid grana, modulate the organization of photosynthetic complexes and significantly alter the dynamics of non-photochemical quenching (NPQ). Taken together, we show that plastidial polyprenols are specialized, environmentally-controlled metabolites that affect the organization of thylakoids and are implicated in the heat stress response of plants.

## Materials and methods

### Plant material and growth conditions

*Arabidopsis thaliana cpt7*, *CPT7-OE*, *aTK* (*hsfa1b,d,e*), *bTK* (*hsfa1a,d,e*), *dTK* (*hsfa1a,b,e*), *eTK* (*hsfa1a,b,d*), *QK* (*hsfa1a,b,d,e*) and *35S:HSFA1A-HA* lines have been previously characterized (Liu et al, 2011; Akhtar et al, 2017; Szaker et al, 2019). For all experiments, seeds were sown on soil and stratified for 3 days. Plants were then grown under standard long day (16 h light/8 h dark) conditions at 22°C, with 70% humidity and 120-150 μmol m-2 s-1 light intensity. For analysis of heat stress responses, plants were heat-treated at 38°C for the indicated time periods and, optionally, recovered at 22°C. For thermotolerance assays (TMHT, BT, ATSR and ATLR) plants were treated as described in Liu et al, 2011 or as indicated on the figure.

### RT-qPCR analysis

For gene expression analysis, rosette leaves of 3-week-old plants were collected. RNA was extracted using a GeneJET RNA Purification Kit (Thermo Scientific), digested with TURBO™ DNase (Ambion) and 1.5 μg were reverse-transcribed using a Transcriptor First Strand cDNA Synthesis Kit (Roche). The obtained cDNA was quantified by qPCR using LightCycler 480 SYBR Green I Master mix (Roche). The level of each transcript was normalized to that of *PROTEIN PHOSPHATASE 2A SUBUNIT A3* (*PP2AA3; At1g13320*). The expression of each gene was examined in three biological replicates. Primers used are listed in Suppl. Table 1.

### Polyprenol extraction and analysis

For polyprenol content analysis, rosette leaves of 3-week-old plants were used. The procedure of polyprenol extraction and analysis was described previously (Akhtar et al, 2017). Briefly, 1.5 g of tissue was freeze-dried, homogenized in a mixture of hexane : acetone [1:1, v/v] using an Ultra-Turrax T25 (IKA Labortechnik), 10 μg of internal standard (Prenol-27; Collection of Polyprenols, Institute of Biochemistry and Biophysics, Polish Academy of Sciences) was added and lipids were extracted for 72 h at room temperature in the darkness. The extracts were filtered and the remaining tissue was re-extracted three times. The extracts were combined, evaporated under a stream of nitrogen, and crude lipids were subjected to alkaline hydrolysis in a mixture of KOH : water : ethanol : toluene 1.14 : 1.14 : 6.2 : 7.5 (w/v/v/v) for 1 h at 95°C. Non-saponifiable lipids were extracted three times with hexane, loaded onto a silica gel 60 column, and purified using isocratic elution with 20% diethyl ether in hexane. Polyprenols were analyzed by HPLC/UV as described previously (Skorupinska-Tudek et al, 2003) using a ZORBAX XDB-C18 (4.6 × 75 mm, 3.5 μm) reverse-phase column (Agilent, USA). Polyprenols were identified relative to authentic standards (Collection of Polyprenols, Institute of Biochemistry and Biophysics, Polish Academy of Sciences). The abundance of polyprenols was examined in three biological replicates.

### Luciferase transactivation assay

To obtain constructs for the luciferase assay, a 700-bp region upstream of the *CPT7* ATG codon (corresponding to the putative *CPT7* promotor) was amplified by PCR, cloned into the pENTR/D-TOPO (Invitrogen) vector and transferred into the *pGWB635* vector (Nakamura et al, 2010) using the LR Clonase II Kit (Invitrogen). Full length *HSFA1A* cDNA was PCR-amplified, cloned into the pENTR/D-TOPO (Invitrogen) vector and subcloned into the *pGWB602* or *pGWB602Ω* vectors (Nakamura et al, 2010) using LR Clonase II Kit (Invitrogen). The resulting *pCPT7:LUC* or *35S:HSFA1A* constructs were transformed into the *Agrobacterium tumefaciens* GV3101 strain. Transformed bacteria were centrifuged, suspended in AS medium (10 mM MES-KOH pH 5.6; 1 mM MgCl_2_), supplemented with acetosyringone (150 μM final concentration) and incubated for 2 h in the dark. Leaves of 1-month-old tobacco (*Nicotiana benthamiana*) plants were infiltrated with transformed Agrobacterium carrying either *pCPT7:LUC* alone or a combination of *pCPT7:LUC* and *35S:HSFA1A*, cultivated for 2-3 days and subjected to a luciferase activity measurement using the NightSHADE LB 985 *in vivo* Plant Imaging System (Berthold). Luciferase activity was examined in discs cut from at least three individual leaves. (Cazzonelli and Velten, 2006)

### Chromatin immunoprecipitation (ChIP)

For ChIP experiments, aerial parts of 3-week-old wild-type and *35S:HSFA1A-HA* plants were collected. Chromatin isolation and ChIP procedure were performed as described by Gendrel et al. (2005). The plant material was frozen in liquid nitrogen, powdered and 2 g were cross-linked. After stopping the cross-linking reaction with glycine (0.125 M final concentration), chromatin was isolated and prepared for ChIP. Anti-HA (Thermo Scientific, 26183) antibodies were coupled to Dynabeads Protein G (Invitrogen) and incubated overnight at 4°C with 10-fold diluted chromatin. After washing, elution, de-crosslinking and digesting with Proteinase K, the obtained DNA was precipitated with isopropanol (1 vol.) and sodium acetate (0.1 vol.), resuspended in water (100 μl) and purified using a QIAquick PCR Purification Kit (QIAGEN). ChIP enrichment of particular gene sequences was determined by qPCR using LightCycler 480 SYBR Green I Master mix (Roche) in three biological replicates.

### Transmission Electron Microscopy (TEM) of chloroplasts

Samples for TEM analysis were prepared as described previously (Kowalewska et al. 2016). Material was cut from the middle part of the fifth leaf of the Arabidopsis rosette. Specimens were analyzed with the help of the JEM 1400 electron microscope (Jeol) equipped with a 11 Megapixel TEM Camera MORADA G2 (EMSIS) in the Nencki Institute of Experimental Biology of the Polish Academy of Sciences, Warsaw, Poland. Ultrastructural grana features such as grana diameter, height, number of layers in particular grana stack, and stacking repeat distance (SRD – distance between adjacent partition gaps in the stack) were calculated with the help of the ImageJ software (Abramoff et al. 2004). Measurements were conducted for 100 grana per each analyzed variant, originating from two biological replicates. Significance was calculated using the one-way ANOVA with post-hoc Tukey test (p ≤ 0.05).

### Spectroscopy measurements of isolated crude thylakoid membrane fraction

For low-temperature fluorescence (77 K), measurements were performed on isolated crude thylakoid membrane fractions diluted to a chlorophyll (Chl) concentration of 10 μg/mL in a 20 mM Hepes-KOH buffer (pH 7.5) containing 330 mM sorbitol, 15 mM NaCl, and 4 mM MgCl2. The crude thylakoid fraction was isolated following the protocol described in Mazur et al, 2016. 77 K fluorescence spectra were recorded using a Shimadzu RF-5301PC spectrofluorimeter in which excitation and emission beams were supplied by optical fibers. Samples containing thylakoids were placed in polytetrafluoroethylene cuvettes and immediately submerged in liquid nitrogen. Excitation wavelength was set at 440 nm and spectra were recorded in the range of 600 to 800 nm.

### Membrane fluidity measurements

Membrane fluidity measurements were performed as described in Bykowski et al, 2021. Briefly, crude thylakoid membrane fractions were diluted to a Chl concentration of 2.5 μg/mL in a 20 mM Hepes-KOH buffer (pH 7.5) containing 330 mM sorbitol, incubated for 10 min at 25°C, supplemented with 1 μM laurdan (6-dodecanoyl-*N*,*N*-dimethyl-2-naphthylamine) and incubated for a subsequent 30 min at 25°C. Steady-state fluorescence emission spectra were recorded at 25°C, followed by heating the sample to 38°C for 5 min and repeating the spectrum measurement. Spectra were recorded in the range of 400–600 nm after excitation at 390 nm using a Shimadzu RF-5301PC fluorimeter. Excitation and emission slits were set to 10 and 5 nm, respectively. Generalized polarization values were calculated according to the formula 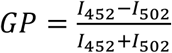, where I_452_ and I_502_ are fluorescence values recorded at wave length 452 nm or 502 nm, respectively.

### Simultaneous measurements of chlorophyll *a* fluorescence and P700 in leaves

*In vivo* fluorescence measurements of 30 min dark-adapted leaves were performed using a Dual-PAM 100 instrument (Heinz Walz GmbH) and photosynthetic parameters were calculated as described in Mazur et al. 2016. Briefly, after determination of minimal fluorescence (F_0_), maximal fluorescence (F_M_) and P700 charge (PM), the actinic light was on (335 μmol photons s^−1^ m^−2^) and leaves were subjected to a series of saturation pulses during four minutes to measure maximum fluorescence (FM’) and P700 charge (PM’). Next, the actinic light was off and during four minutes of dark-recovery phase FM’ and PM’ values were measured again. Alternatively, light induction curves with low intensity actinic light (50 μmol photons s^−1^ m^−2^) were recorded in the same manner as described before, except of the dark-recovery phase. Next, the light curve was recorded during illumination of light adapted leaves by actinic light with exponentially increasing intensity (from 5 to 5000 μmol photons s^−1^ m^−2^). The measurements were conducted for three biological replicates.

### Preparation of thylakoid-mimicking lipid multibilayers

MGDG and DGDG lipids were purchased from Larodan. *n*-Dodecyl-β-D-maltoside (DM), Tricine and KCl were obtained from Sigma-Aldrich. Mixture of polyprenols (Pren-9, −10 and −11) was provided by the Collection of Polyprenols, Institute of Biochemistry and Biophysics, Polish Academy of Sciences. Light harvesting complexes LHCII were isolated from fresh spinach (*Spinacia oleracea L*.) leaves as described previously (Krupa et al, 1987). LHCII-multibilayers were prepared as previously (Janik et al, 2013). Briefly, MGDG and DGDG were dissolved in chloroform at a molar ratio of 2:1. The LHCII complexes were suspended in 20 mM Tricine and 10 mM KCl buffer, pH 7.6, containing 0.03% DM. The mixture of MGDG and DGDG was transferred to glass test tubes and placed in a vacuum (<10^−5^ bar) for 30 min. Polyprenol was dissolved in hexane (5 mg in 0.5 ml) and, when indicated, added at the beginning of evaporation to the MGDG and DGDG mixture at a molar fraction of 0.5% or 5%. LHCII complexes were transferred to lipid films and incorporated into lipid membranes through mild sonication in an ultrasonic bath for 30 min. The molar ratio of LHCII complexes to lipids was 1:200. In order to remove DM, the samples were incubated with Bio-beads adsorbent at 4°C for 12 h. Multilayers of LHCII and lipids were separated from the samples by centrifugation at 15,000 *g* for 5 min. The pellets were resuspended in 20 mM Tricine and 10 mM KCl buffer and used for FLIM measurements.

### Fluorescence Lifetime Imaging

Fluorescence Lifetime Imaging analyses were carried out on a confocal MicroTime 200 (PicoQuant GmbH, Germany) system coupled to an inverted microscope OLYMPUS IX71. The samples were illuminated with a 470 nm pulse laser with a repetition rate of 10 MHz. The lifetime resolution was better than 16 ps. Photons were collected with a 60x water-immersed objective (NA 1.2, OLYMPUS UPlanSApo). A single focal plane was selected with a pinhole diameter of 50 μm. Scattered light was removed using a dichroic ZT473RDC XT (ANALYSENTECHNIK), 470 notch filter (Chroma Technology) and observations were performed with the addition of a 690/70 band pass filter. Results of the measurements were analyzed using the SymPhoTime 64 software (PicoQuant GmbH, Germany).

### Profiling of lipids from thylakoid membranes

Total lipids were extracted from isolated *Arabidopsis thaliana* thylakoid membranes (an amount of sample yielding 100 μg of chlorophyll *a*) by addition of chloroform : methanol (2:1, v/v) to obtain a final concentration ratio of chloroform : methanol : homogenate 8:4:3 (v/v) according to Folch et al, 1957. After phase separation, the organic phase was collected and washed with 5 M NaCl. The lipid-containing organic extracts were dried under a gentle stream of nitrogen, the obtained oily residue was dissolved in chloroform : methanol (2:1, v/v) and stored at −20 °C until lipid determination by high performance thin layer chromatography (HPTLC) was performed.

Lipid separation was performed as described before (Schaller et al, 2010). Briefly, concentrated lipid extracts with a defined chlorophyll *a* concentration (in a range of 5-10 μg) were applied to HPTLC silica gel 60 plates (Merck, Germany) with the CAMAG Linomat 5 in the form of bands and developed in TLC chambers based on a method described in Ventrella et al, 2007 with chloroform : methanol : acetic acid : water (75 : 13 : 9 : 3, v/v) as eluent. After the HPTLC separation, the lipids were stained with primuline (0.05% in acetone : water (80:20, v/v)) according to White et al, 1998. A TLC Visualizer CAMAG (Muttenz, Switzerland)) in UV 366 nm mode was used to visualize the HPTLC plates and the spots were analyzed using the software package WinCat provided by CAMAG (Muttenz, Switzerland). The content of the particular species of lipids was calculated in relation to lipid standards, 2.5 μg each (Lipid Products, South Nutfield, UK), run on the same plate. The results were expressed relative to the amount of chlorophyll *a*.

### Glycoprotein staining

Protein extracts from chloroplasts (25 μg of protein per line) were separated by SDS-PAGE and transferred to nitrocellulose membrane. The membranes were probed for 16 h with Concanavalin A conjugated with horseradish peroxidase (1 mg/mL; Sigma-Aldrich), washed for 45 min with PBS, and incubated for 30 s with SuperSignal West Pico Chemiluminescent Substrate (Thermo Scientific). Detection of chemiluminescence signal was performed with Molecular Imager ChemiDoc XRS+ (Bio-Rad).

### HILIC-HPLC analysis of chloroplast protein-derived *N*-glycans

For glycan analysis, chloroplasts were isolated from 5 g of Arabidopsis leaves according to the manual (Weigel and Glazebrook, 2002). The chloroplast proteins were de-glycosylated and fluorescently labeled as described previously (Link-Lenczowski et al., 2018). Briefly, protein pellets were dissolved in 20 μl of PNGase F denaturation buffer (New England Biolabs, Ipswich, MA) and denatured at 95°C for 10 min. Subsequently, 20 μl of digestion buffer were added to the samples, and proteins were de-glycosylated with PNGase F (500,000 U/ml, New England Biolabs) overnight at 37°C. The released *N*-glycans were purified on graphitized carbon SPE columns (Supelclean™ ENVI-Carb™, Sigma-Aldrich, Poznań, Poland) and labeled with a fluorescent tag (anthranilic acid, 2-AA). 2-AA-labeled N-glycans were purified on SPE columns (Spe-ed Amide-2, Applied Separations, Allentown, PA), eluted with HPLC-grade water and stored at −20°C for further analysis.

The obtained glycans were analyzed by HILIC-HPLC on a TSK gel-Amide 80 column (4.6×150 mm, 3 μm bead size, Tosoh Bioscience) with the Shimadzu Prominence HPLC system (Shimadzu, Duisburg, Germany) and an in-line RF-20Axs fluorescence detector as described previously with some modifications (Link-Lenczowski et al., 2019). Solvent A was acetonitrile and solvent B was 50 mM ammonium hydroxide, titrated to pH 4.4 with formic acid, in HPLC-grade water. Samples were loaded in 70% acetonitrile and chromatography was performed at 30°C using gradient conditions as follows: 0 min, 35% B; 22 min, 46% B; 23 min, 100% B; 25 min, 100% B; 27 min, 35% B; 35 min, 35% B at a solvent flow rate of 1 mL/min. Each sample set was run together with the external standard (2-AA labeled dextran ladder, Glyko Prozyme, Hayward, CA). In each chromatogram 15 individual peaks were identified and peak assignments were made basing on glucose units (GU), determined from a standard. For quantitative analysis, the total area of all glycan-peaks was normalized to the protein concentration. The results are presented as mean values and differences between groups were tested for significance with the two-tailed Student’s t-test using the GraphPad Prism trial version (GraphPad Software, La Jolla, CA). Analysis was performed in triplicate.

### MALDI-ToF-MS analysis of chloroplast-derived *N*-glycans

2-AA-labeled *N*-glycans prepared for HILIC-HPLC were desalted on C18 ZipTips (Millipore, Merck, Warszawa, Poland), following the manufacturer’s instructions. Purified N-glycans were eluted directly on MALDI stainless steel target plates along with 2,5-dihydroxybenzoic acid solution (10 mg/ml in acetonitrile : water (1:1, v:v) containing 0.1% TFA) and left to dry under atmospheric pressure at room temperature. MALDI-ToF-MS was performed on an UltrafleXtreme™ mass spectrometer (Bruker Daltonics, Bremen, Germany). The external calibration of the instrument was performed with the use of the Bruker peptides calibration kit (757.4 Da – 3147.5 Da). The mass spectra were acquired in the negative ion reflectron mode over the *m/z* range from 700 to 5000 for a total of 5000 shots/sample spot. *N*-glycan structures were identified basing on the *m/z* values and on common knowledge of glycobiology.

## Results

### Expression of *CPT7* and polyprenol biosynthesis are upregulated in heat stress

Recent studies revealed that CPT7 is responsible for the biosynthesis of polyprenols (Pren-9, −10 and −11) in Arabidopsis (Akhtar et al, 2017). However, molecular factors regulating the expression of *CPT7* and the synthesis of polyprenols remain unknown. While examining our transcriptomic data (Buszewicz et al, 2016) and publicly available microarray databases (Arabidopsis eFP Browser at http://bar.utoronto.ca; Winter et al, 2007) we have found that heat stress upregulates the expression of *CPT7* (Suppl. Fig. 1A). We have confirmed this observation by RT-qPCR analysis of *CPT7* mRNA levels in 3-week-old plants exposed to heat (38°C) for 3 hours (Suppl. Fig. S1B). The level of *CPT7* mRNA started to increase as soon as 15 min after the beginning of treatment, reached its peak after 1h and then stared to decline. Plants grown in parallel in control conditions showed no significant changes in *CPT7* transcript level (Suppl. Fig. S1B), indicating that the upregulation of *CPT7* was the result of heat treatment and was not related to the diurnal cycle. Moreover, *CPT7* was the only *cis*-prenyltransferase in Arabidopsis leaves whose expression was substantially affected during heat treatment (Suppl. Fig. 2), suggesting that *CPT7* might be specifically involved in the response to heat stress.

To investigate in detail the induction of *CPT7* transcript upon heat stress and its impact on CPT7-derived poliprenol content, we applied long-term exposure to high temperature (24 h in 38°C) and measured *CPT7* mRNA and polyprenol content in multiple time points across the experiment. In accordance with previous results, *CPT7* transcript level reached a peak of expression at 1 h of heat treatment, followed by a rapid decrease to levels similar to those observed in non-stressed plants (Fig. 1A). However, quantification of CPT7-derived polyprenols showed that increase of their content starts at approx. 3 h when *CPT7* mRNA level is already declining and continues as the heat treatment persists (Fig. 1B), demonstrating that exposure to high temperature causes accumulation of polyprenols. Interestingly, polyprenol level remained elevated during recovery after heat stress, although it displayed small decrease over time which is opposite to the trend described in non-stressed plants which accumulate polyprenols as they age (Fig. 1C) (Surmacz and Swiezewska, 2011). This opens the possibility that polyprenols might undergo catabolic degradation in plant tissues by a so far uncharacterized mechanism. To check whether any of the remaining CPTs can substitute CPT7 in heat-induced production of the analyzed polyprenols, we examined their content in a heat stressed *cpt7* null mutant. This analysis revealed that polyprenols-9, −10 and −11 are missing in both stressed and non-stressed *cpt7* plants (Suppl. Fig. 3), confirming previous report (Akhtar et al, 2017) that CPT7 is the only enzyme responsible for the biosynthesis of these polyprenols and showing that other *cis*-prenyltransferases cannot compensate for its deficiency also in heat stress conditions.

**Figure 1.**
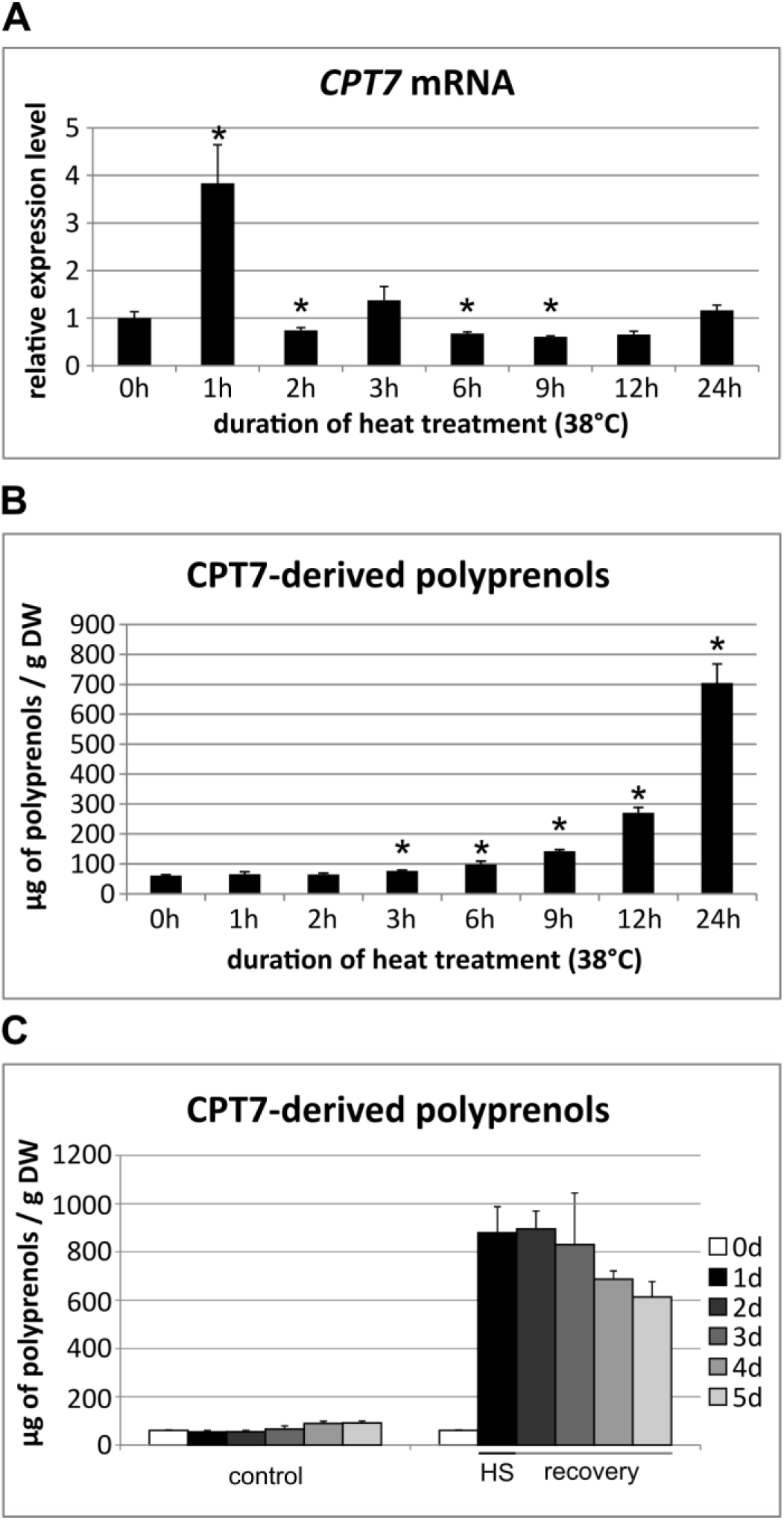
Heat stress induces *CPT7* expression and accumulation of CPT7-derived polyprenols in Arabidopsis leaves. **A)** RT-qPCR analysis of mRNA levels of *CPT7* during heat treatment. Expression levels are relative to *PP2AA3* and further normalized to 0 h. **B)** Polyprenol content during heat treatment. **C)** Polyprenol content in control (non-stressed, 0d) plants or during heat treatment (HS) and recovery. For all experiments means and SDs from three biological replicates are shown. Statistical relevance of the results was checked using the t-test, always in relation to the 0 h time point; * – P-value < 0.05.

Taken together, we have found that heat stress causes a rapid and transient induction of CPT7 expression that is followed by accumulation of CPT7-derived polyprenols Pren-9, −10 and −11. The level of polyprenols remains increased for days after heat treatment is over.

### HSFA1 transcription factors directly regulate the expression of *CPT7* and accumulation of polyprenols in heat stress

Heat shock transcription factors (HSFs) are master regulators of the plant heat stress response, acting as activators or repressors of genes involved in acclimation to high temperature (Scharf et al, 2012). As *CPT7* is upregulated shortly after applying heat stress, we decided to check whether transcription factors of the HSFA1 group, acting at early stages of the heat response (Liu et al, 2011; Yoshida et al, 2011), are responsible for the activation of *CPT7* expression. As all four HSFA1s are highly redundant, we used quadruple (*QK*) *hsfa1* mutants (Liu et al, 2011) and measured *CPT7* mRNA level before and during heat treatment. RT-qPCR analysis showed that, in contrast to WT lines, heat treatment resulted in only minor upregulation of the level of *CPT7* transcript in *QK* plants (Fig. 2A). This indicated that HSFA1s are required for full induction of *CPT7* expression. To check if individual HSFA1s are sufficient for the induction of *CPT7*, we measured the level of *CPT7* mRNA in triple (*TK*) *hsfa1* mutant lines (Liu et al, 2011) before and after 1 h of heat treatment. As expected, heat-treated *bTK* (*hsfa1a,d,e*), *dTK* (*hsfa1a,b,e*) and *eTK* (*hsfa1a,b,d*) lines displayed downregulation of the *CPT7* transcript (Fig. 2C), resulting in an intermediate level of transcript when compared to WT and *QK* lines, showing that HSFA1B, HSFA1D and HSFA1E are individually partially able to induce *CPT7* transcription. On the other hand, HSFA1A alone was sufficient to activate the expression of *CPT7* since the *CPT7* mRNA was not downregulated in *aTK* (*hsfa1b,d,e*) plants (Fig. 2C). Interestingly, the *CPT7* transcript level was also decreased in non-stressed *QK* plants when compared to the WT line (Fig. 2B), indicating that HSFA1s regulate its expression not only upon heat stress but also in standard growth conditions.

**Figure 2.**
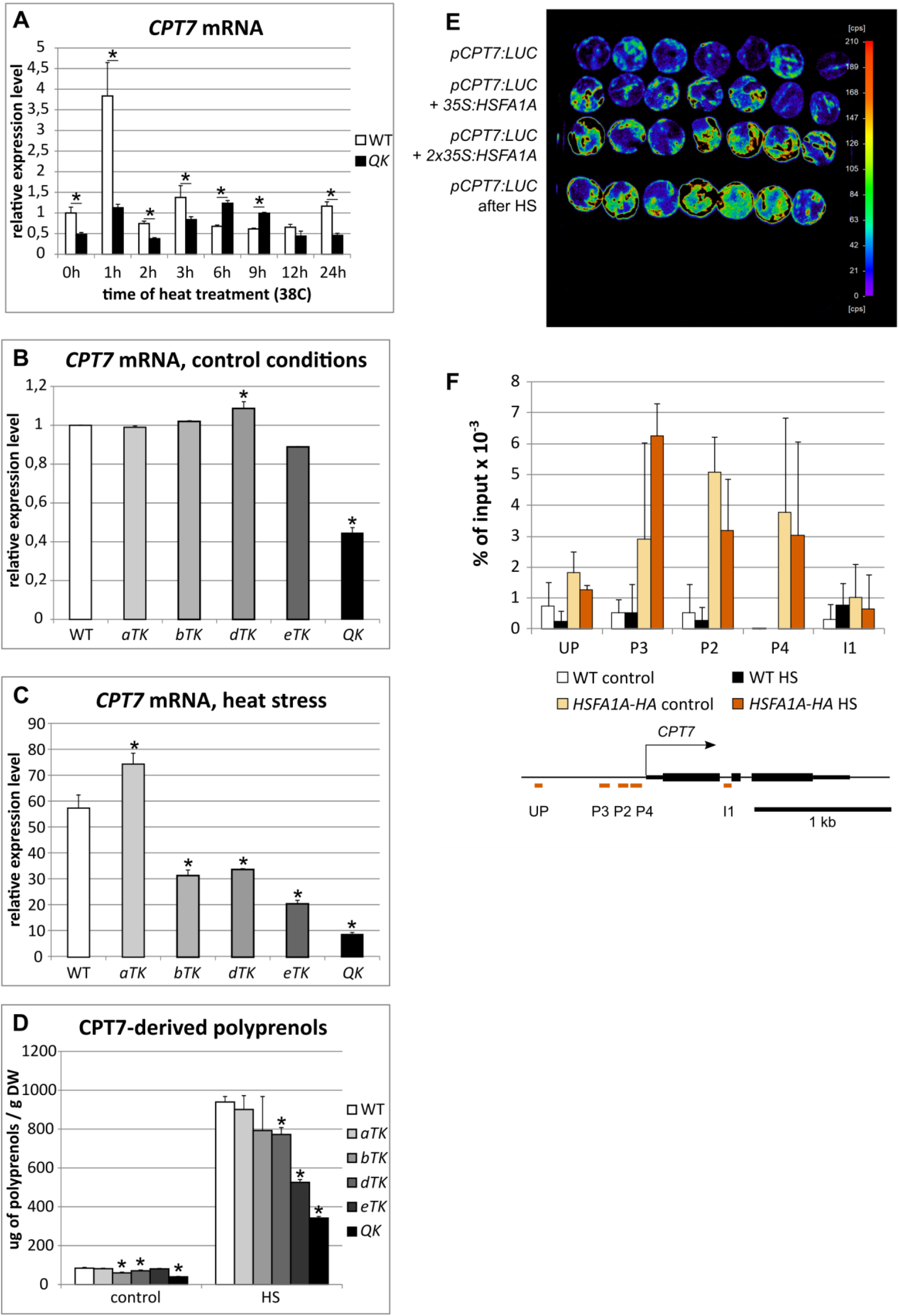
HSFA1 transcription factors regulate the expression of *CPT7* and polyprenol accumulation in both control and heat stress conditions. **A)** RT-qPCR analysis of mRNA levels of *CPT7* in wild type and *QK* plants during heat treatment. Expression levels are relative to *PP2AA3* and further normalized to 0 h in WT. **B, C)** RT-qPCR analysis of mRNA levels of *CPT7* in wild type and *hsfa1* triple (*TK*) and quadruple (*QK*) mutant plants before and after 1 h of heat treatment. Expression levels are relative to *PP2AA3* and further normalized to 0 h in WT. **D)** Quantitative analysis of polyprenol content in wild type and *hsfa1* triple (*TK*) and quadruple (*QK*) mutant plants grown in standard conditions or after 24 h of heat treatment. **E)** HSFA1A and heat activate the expression of a *pCPT7:LUC* reporter in tobacco leaves. Shown is the light intensity of the luciferase (LUC) reporter expressed in tobacco leaves either alone or together with the HSFA1A trans-activator. The results of a representative experiment are shown. **F)** HSFA1A-HA occupancy at the *CPT7* locus assayed by ChIP-qPCR, before and after 1h of heat treatment (38°C). Positions of the analyzed regions: upstream of the promoter (UP), in the promoter (P2-4) and in the first intron (I1) of the *CPT7* DNA sequence, are indicated below. Data are % of input values. For all experiments means and SDs from three biological replicates are shown.. Statistical relevance of the results was checked using the t-test, always in relation to WT; * – P-value < 0.05.

Next we checked if the lack of HSFA1s affects the accumulation of polyprenols. Consistent with the RT-qPCR analysis, the total amount of polyprenols was significantly decreased in both stressed and non-stressed *QK* plants when compared to WT (Fig. 2D). Moreover, after heat stress, *TK* plants contained intermediate levels of polyprenols, relative to WT and *QK*. Again, these results show that HSFA1 transcription factors are partially redundant in the regulation of *CPT7* activity and that they are necessary for its proper function.

To further test the ability of HSFA1A to activate the expression of *CPT7*, we performed a transactivation assay using luciferase as a reporter. Tobacco leaves were transiently co-transformed with *pCPT7:LUC* and *35S:HSFA1A* constructs and the luminescence level of LUC was measured. The leaves co-transformed with both constructs displayed higher light intensity, compared to control leaves transformed only with *pCPT7:LUC* (Fig. 2E), showing that HSFA1A is able to activate the *CPT7* promoter. Additionally, heat treatment of the leaves transformed only with the reporter construct also increased the activity of luciferase, confirming that the *CPT7* promoter is responsive to heat stress, possibly with involvement of endogenous tobacco HSFs (Sanmiya et al, 2020) acting as expression activators.

Binding of HSF transcription factors to chromatin relies on the presence of HSE (Heat Shock responsive Element) motifs in the DNA sequence (Pelham, 1982; Amin et al, 1988). While the putative *CTP7* promoter used in the transactivation assays does not contain classical HSEs, two non-canonical HSF binding sites (TTCCAACCTTC and CCCCT; Guo et al, 2008) were identified upstream of the start codon. To verify if HSFA1A binds to the *CPT7* promoter and directly regulates *CPT7* expression, a ChIP experiment using a *35S:HSFA1A-HA* line was performed. Quantitative analysis of DNA precipitated with the anti-HA antibody showed a significant enrichment in DNA sequences, comprising both non-canonical HSE sites, located upstream of the start codon of the *CPT7* gene (Fig. 2F).

Taken together, the results of genetic, biochemical and molecular experiments described above clearly indicate that expression of *CPT7* and consequently the biosynthetic route of polyprenols are directly regulated by HSFA1s, in particular by the HSFA1A transcription factor.

### CPT7-derived polyprenols affect biochemical features of chloroplasts

Recent studies have shown that polyprenols, Pren-9, −10 and −11, are components of the thylakoid membrane system (Akhtar et al, 2017). Therefore, we checked if disturbances in the level of these highly unsaturated molecules cause any further changes in the lipid composition of thylakoid membranes. To this end, we quantified the lipid content of thylakoids isolated from WT, *cpt7* and *CPT7-OE* plants grown in either control or heat stress conditions (Fig. 3A). For all analyzed genotypes, HPTLC analysis revealed the presence of four major thylakoid membrane lipid species: monogalactosyldiacylglycerols (MGDG), digalactosyldiacylglycerols (DGDG), sulfoquinovosyldiacylglycerols (SQDG) and phosphatidylglycerols (PG). Since we could not detect polyprenols using this technique, we analyzed samples from the same thylakoid preparation by HPLC-UV for polyprenol quantification. As expected, the dominant lipids in all studied samples were galactosyldiacylglycerols (MGDG and DGDG). Interestingly, the content of polyprenols was comparable to that of SQDG, showing that polyprenols should be considered major constituents of thylakoid membranes. As expected, we could not detect CPT7-derived polyprenols in the *cpt7* mutant while their content in *CPT7-OE* plants was doubled compared to WT. Interestingly, *cpt7* plants grown in control conditions displayed a lowered level of MGDG, while overproduction of polyprenols in *CPT7-OE* plants did not cause any significant changes in the content of other lipid classes. There are multiple studies showing that heat stress affects the lipid composition of thylakoid membranes (Tang et al, 2016; Higashi et al, 2018; Shiva et al, 2020). In our analysis, heat treatment caused a substantial increase in the absolute content of MGDG and DGDG across all studied genotypes and inflated the difference in MGDG content between WT and *cpt7* plants (Fig. 3A). As expected, similarly to studies done on entire leaves, heat stress caused an increase in the level of polyprenols in thylakoids both in WT and *CPT7-OE* plants (Fig. 3A).

**Figure 3.**
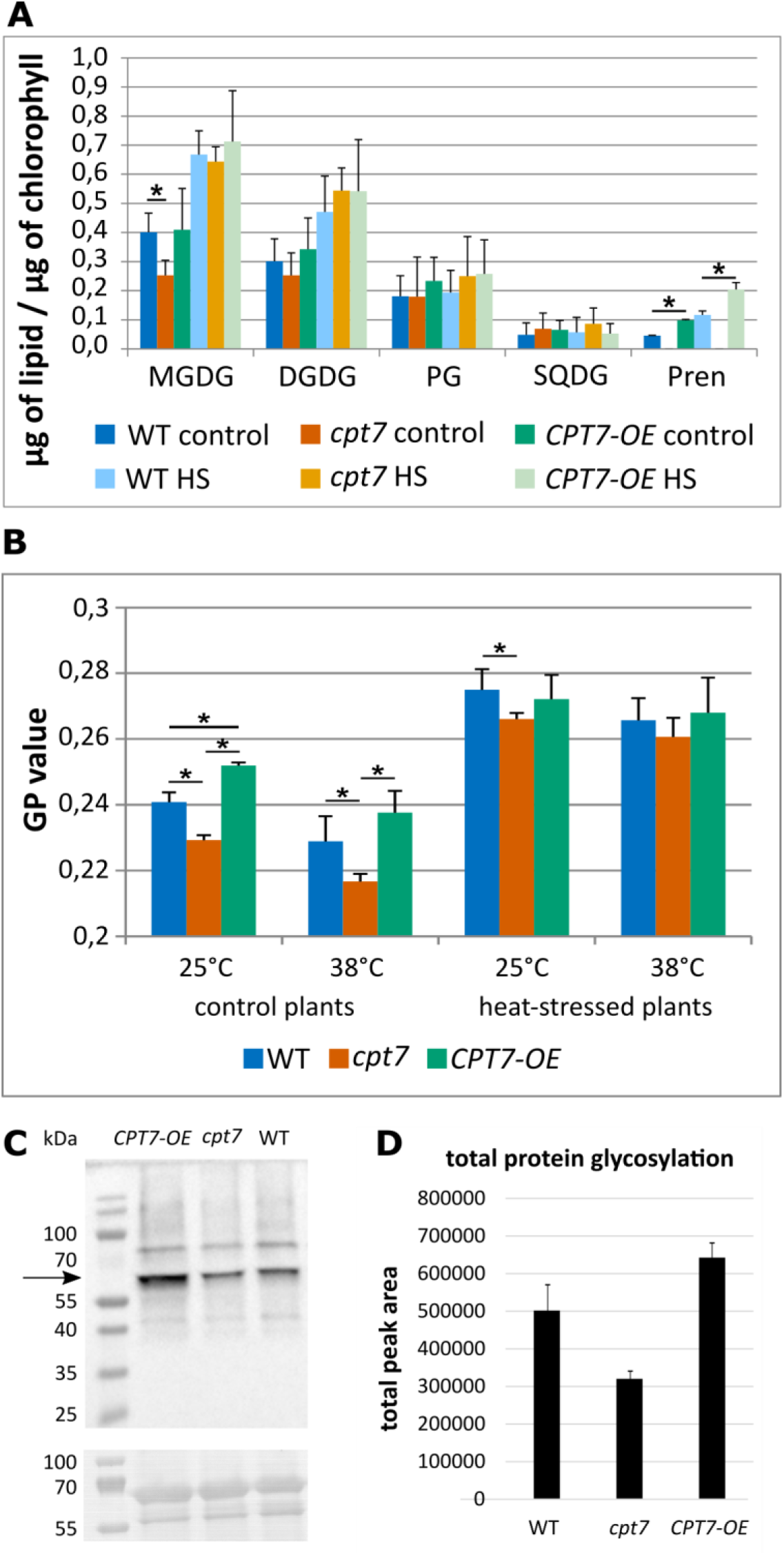
Protein glycosylation, lipid composition and fluidity of membranes are affected by polyprenols. For protein glycosylation analysis, proteins isolated from chloroplasts of WT, *cpt7* or *CPT7-OE* plants were used. **A)** Content of dominating lipid constituents of thylakoid membranes. **B)** Membrane fluidity measurement employing Laurdan as a reporter molecule is expressed as generalized polarization (GP), see Materials and Methods section. Individual thylakoid samples were analyzed in both 25°C and 38°C. **C)** Concanavalin A staining of total chloroplast protein extracts (upper panel); Ponceau S staining (bottom panel) was used as a loading control. **D)** Total content of glycan moieties released from chloroplast proteins. The sum of peak areas corresponding to identified glycans was normalized to protein content. Means and SDs from three biological replicates are shown. For the analyses of lipid profile and membrane fluidity thylakoids of WT, *cpt7* and *CPT7-OE* plants grown in standard or heat stress (24h at 38°C) conditions were isolated: For all experiments means and SDs from three biological replicates are shown. Statistical significance was checked using the t-test; * – P-value < 0.05.

Previous studies done on Arabidopsis (Akhtar et al, 2017) and tomato (*Solanum lycopersicum*) plants (Van Gelder et al, 2018) have shown that polyprenols reduce the fluidity of thylakoid membranes. Since the plants used in both studies were grown in control conditions, we decided to check if heat treatment further modulates the effect of polyprenols. To this end, we employed Laurdan as a reporter to measure membrane fluidity of thylakoids isolated from WT, *cpt7* and *CPT7-OE* plants grown in either standard or heat-stress conditions. Moreover, all thylakoid samples were analyzed in both 25°C and 38°C to check if polyprenol levels can affect changes in fluidity caused by elevated temperature. In accordance to previous reports, a lack of polyprenols resulted in a more fluid state of thylakoid membranes (expressed by a lower GP value) obtained from plants grown in standard conditions (Fig.3B). Heating of the thylakoid samples to 38°C resulted in higher membrane fluidity across all genotypes, although the effect of polyprenols remained unchanged. When analyzing heat-stressed plants, a substantial increase in membrane rigidness could be observed for all plant lines, which could be attributed to acclimatization to high temperature. Interestingly, the effect of polyprenol deficiency became much less pronounced in heat-stressed plants. These results indicate that while polyprenols are heat-responsive lipids clearly affecting membrane fluidity, their main biological function might not be to counteract the liquefying effect of high temperature exerted on thylakoid membranes.

Moreover, further characterization of *cpt7* and *CPT7-OE* plants revealed that disturbances in polyprenol levels affect the glycosylation status of chloroplast proteins. Staining of chloroplast protein extracts with the carbohydrate-binding lectin Concanavalin A revealed a protein band migrating at approx. 60 kDa that apparently produced a stronger signal in lines containing higher levels of polyprenols (*CPT7-OE*) than in the WT line (Fig. 3C). Conversely, this signal was weaker in the *cpt7* mutant. To confirm this observation, glycan moieties were excised from chloroplast proteins and analyzed using HPLC. Structural characterization of the released *N*-glycans with MALDI-ToF mass spectrometry revealed the presence of 21 individual glycan species (Suppl. Table 2). The major signal was detected at *m/z* 1760.6 – it corresponds to a diantennary complex structure with terminal galactose residues (Hex5HexNAc4-AA; Suppl. Fig. 4). Additionally, structures with a single xylose residue, characteristic for plant glycans (Kaneko et al, 2016), were detected at *m/z* 1162.3 (Hex3HexNAc2Pen1-AA) and *m/z* 1365.4 (Hex3HexNAc3Pen1-AA). Although the glycomic profiles of WT, *cpt7* and *CPT7-OE* lines were similar (Suppl. Fig. 5; Suppl. Table 2), quantitative differences between the tested lines were noted. In accordance with previous results, we have found a decrease and increase in the total glycosylation level in *cpt7* and *CPT7-OE* plants, respectively (Fig. 3D). These results suggest that polyprenols might be implicated, most likely indirectly, in the glycosylation of chloroplast proteins.

### Polyprenols affect the ultrastructure of the chloroplast thylakoid network

Since heat stress is known to affect photosynthesis (Hu et al 2020) and polyprenols are plastidial lipids accumulating in response to heat treatment, we performed in-depth studies on how polyprenols affect structural and functional aspects of the chloroplast thylakoid network. Previous studies have shown that deficiency of plastidial polyprenols affects photosynthetic performance in Arabidopsis (Akhtar et al, 2017). Therefore, we decided to check if polyprenols affect the ultrastructure of chloroplasts and influence heat-induced changes in their organization. The ultrastructure of mesophyll chloroplasts of all examined genotypes in both control and heat stress conditions was typical for Arabidopsis plants (Suppl. Fig. 6). The thylakoid network of all examined plants was formed by stacked grana thylakoids and unstacked stroma thylakoids connected tightly to each other (Fig. 4 A-F). To address detailed ultrastructural differences between the examined plants, measurements of grana structural parameters were performed (Fig. 4 G-J). A significantly lower diameter was measured for heat-stressed *cpt7* plants when compared to WT (Fig. 4 G). Moreover, the height of grana stacks was lower in *cpt7* plants when grown in control conditions (Fig. 4 H, I). The most noticeable differences were registered for the stacking repeat distance (SRD) values of the analyzed grana (Fig. 4 J). The SRD parameter corresponds to the thickness of a single thylakoid (membrane – lumen – membrane) in the grana stack together with the neighboring partition gap and therefore reflects inversely the degree of membrane stacking. Heat-stress conditions induced an increase in grana compactness in all examined plants resulting in a decrease of the mean SRD value of about 2 nm for WT and *CPT7-OE* plants and below 1 nm for *cpt7* plants. The decrease in the SRD value observed for heat-stressed *cpt7* plants was smaller than for the other two genotypes, most likely due to the lower initial SRD value registered for *cpt7* control plants. In control temperature conditions, the SRD parameter values correlated with the content of thylakoid polyprenols - the higher the content, the lower was the degree of grana stacking.

**Figure 4.**
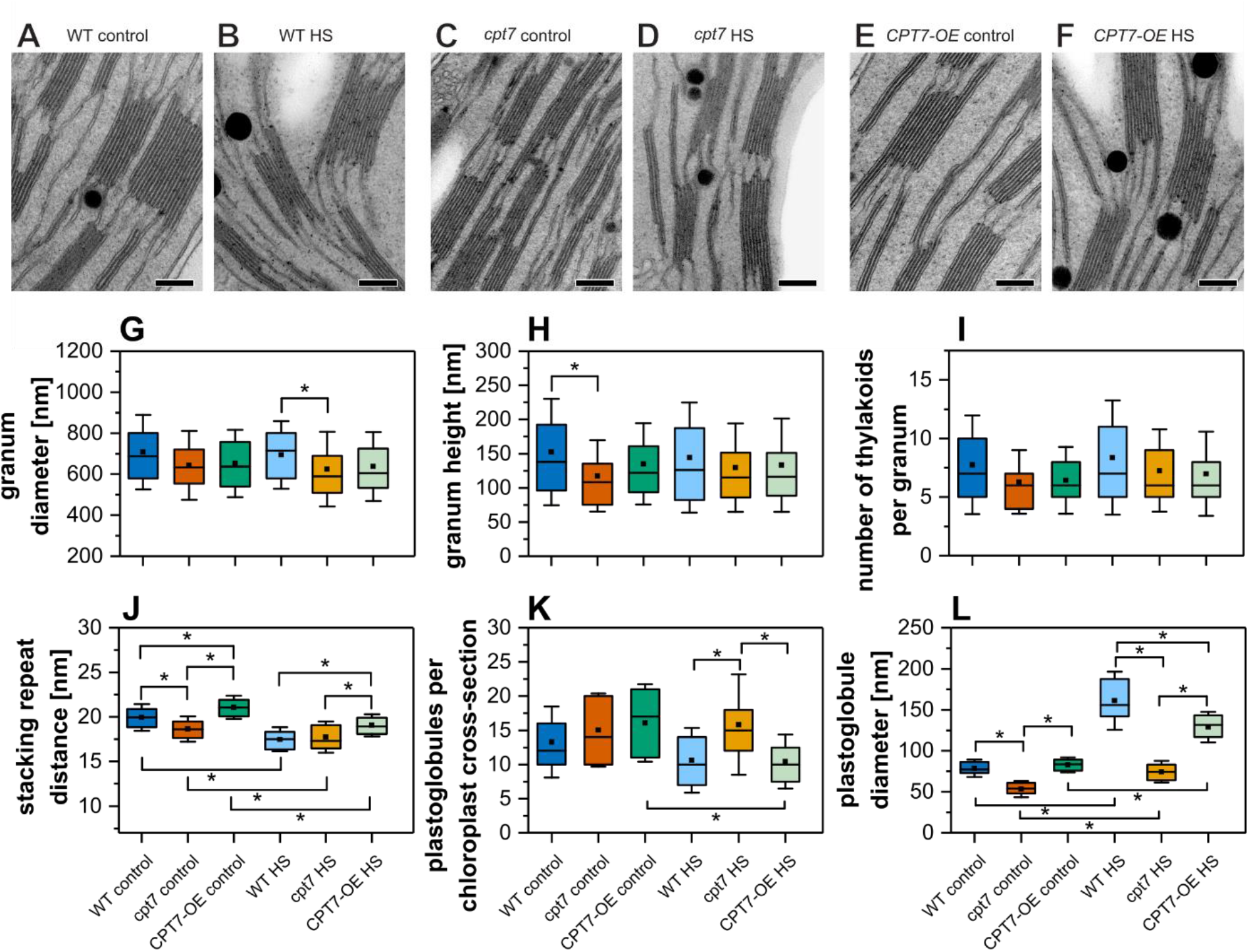
Polyprenols affect the structure of the thylakoid network of Arabidopsis plants. Thylakoids were isolated from WT, *cpt7* or *CPT7-OE* plants grown in standard or heat stress (24h at 38°C) conditions. **A-F** – electron micrographs of mesophyll chloroplasts of WT (**A, B**), *cpt7* (**C, D**) and *CPT7-OE* (**E, F**) plants in control (**A, C, E**) and HS conditions (**B, D, F**). **G-L** – grana structural parameters: diameter (**G**), height (**H**), number of thylakoids per granum (**I**), stacking repeat distance (SRD) (**J**), number of plastoglobules per chloroplast cross-section (**K**) and plastoglobule diameter (**L**). Observations were performed for three independent biological replicates. Numeric data are means and SDs from at least 25 independent measurements. Statistical relevance of the results was checked using one-way ANOVA with post-hoc Tukey test; * – P-value < 0.05.

Studies performed on tomato plants showed that polyprenol deficiency increases plastoglobule number per chloroplast (Van Gelder et al, 2018). While we have not found changes in plastoglobule count between the tested Arabidopsis lines when grown in control conditions, there was a significant increase in the number of plastoglobules in the heat-stressed *cpt7* line when compared to both WT and *CPT7-OE* plants (Fig. 4K). Moreover, there was a significant decrease in the plastoglobule diameter in *cpt7* plants grown in control conditions (Fig. 4L). When heat stress was applied, an increase in plastoglobule diameter was observed across all lines, although it was significantly lower for *cpt7*.

### Polyprenols modulate arrangement of the photosynthetic apparatus

Previous studies have shown that lack or a decreased content of polyprenols cause a decrease in the linear electron transport rate, resulting in a lowered quantum yield of photosystem II (Y(II) (Akhtar et al, 2017). However, changes in the ultrastructure of chloroplasts in *cpt7* and *CPT7-OE* plants suggest that polyprenols might also affect arrangement of the photosynthetic complexes. Therefore, we employed a low-temperature (77 K) chlorophyll fluorescence measurement technique that can assess changes in the organization of photosynthetic complexes. The emission spectra obtained for WT, *cpt7* and *CPT7-OE* plants, grown in control or heat stress conditions (Fig. 5A), had a shape typical for Arabidopsis plants (Mazur et al. 2019). The difference spectra revealed that both the *cpt7* mutant and the *CPT7-OE* line displayed increased aggregation of LHCII complexes (positive peak at 700 nm) and altered contribution of photosystem I (PSI, peak at 734 nm) and photosystem II (PSII, peak at 682 nm) (Fig. 5B). Moreover, a higher contribution of photosystem II and a lower contribution of photosystem I in plants subjected to heat stress were observed for all analyzed genotypes, with the highest increase of PSII contribution observed for *cpt7* (Fig. 5C). These results show that heat treatment affects the organization of chlorophyll-containing protein complexes and that their organization can be further affected by polyprenol content.

**Figure 5.**
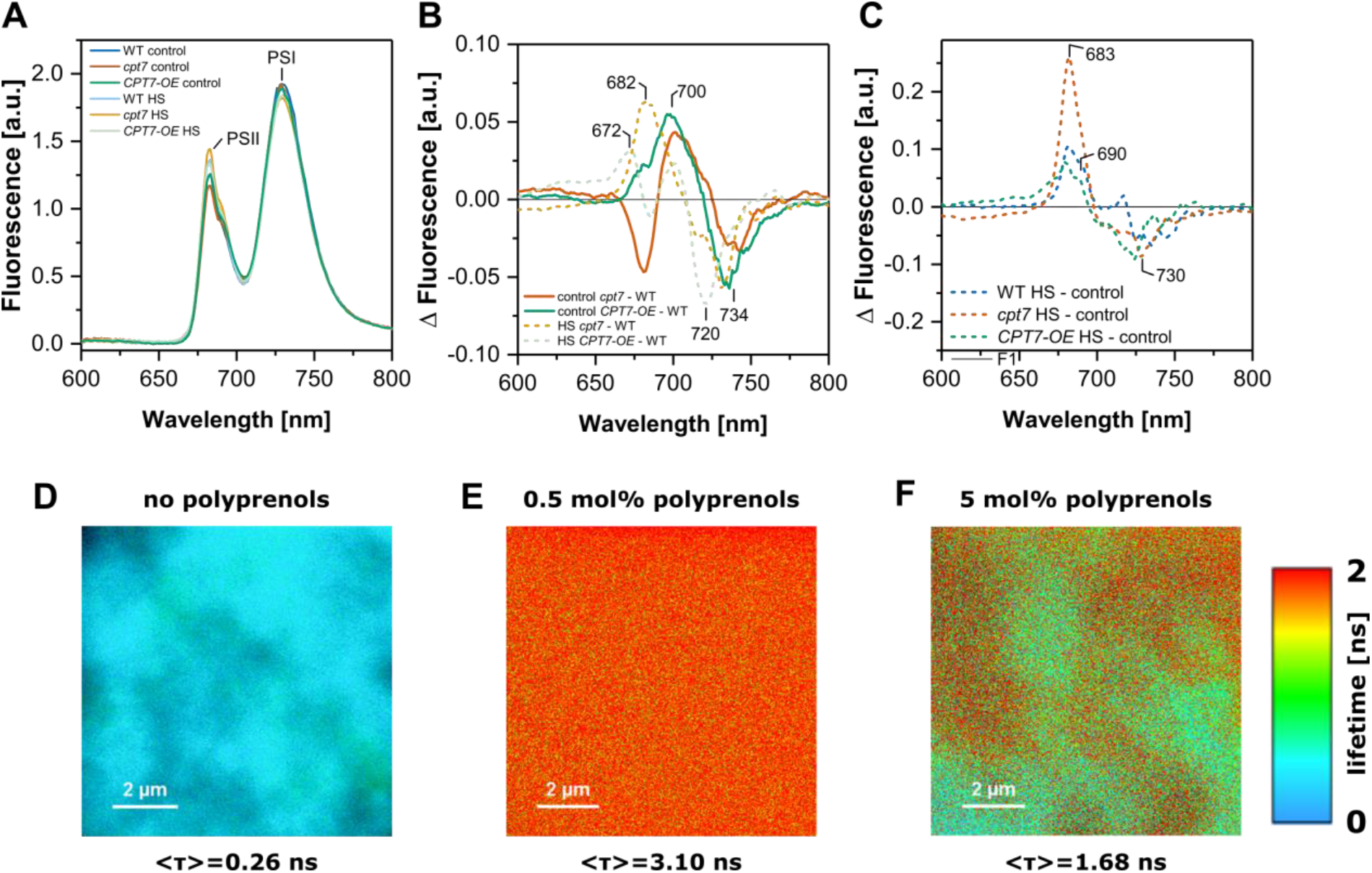
Polyprenols affect the molecular organization of photosynthetic complexes. **A-C** – low temperature (77 K) fluorescence emission spectra (ex 440 nm) of control and HS treated plants (**A**); differential *cpt7*-minus-WT and *CPT7-OE*-minus-WT fluorescence emission spectra in control and HS conditions (**B**); differential HS-minus-control fluorescence emission spectra in WT, *cpt7* and *CPT7-OE* plants (**C**); fluorescence spectra were normalized to an equal area under the curves. Chloroplasts isolated from leaves of WT, *cpt7* and *CPT7-OE* Arabidopsis plants grown in standard or heat stress (24 h at 38°C) conditions were analyzed. **D**, **E**, **F**) FLIM images of fragments of lipid multibilayers containing LHCII without additions (**D**) or supplemented with 0.5 mol% (**E**) or 5 mol% (**F**) of polyprenols. Amplitude-averaged fluorescence lifetimes, calculated over entire images, are presented below each panel. Representative images are shown.

To further study the effects of polyprenols on the organization of the photosynthetic apparatus, we employed fluorescence lifetime imaging microscopy (FLIM) analysis of thylakoid stack-mimicking multibilayers formed from a mixture of chloroplast lipids (MGDG:DGDG, 2:1, mol/mol) and functional pigment-protein antenna complexes LHCII. In accordance with previous results (Janik et al, 2013), the relatively short average fluorescence lifetime of chlorophyll *a* in such a system indicates aggregation of LHCII, which results in excitation quenching (Fig. 5D). An addition of polyprenols to the lipid phase (0.5 mol%) resulted in a pronounced increase in the average fluorescence lifetimes (Fig. 5E). Interestingly, the effect of polyprenols weakened when they were applied in a higher concentration (5 mol%), although their de-aggregating effect was still apparent (Fig. 5F). These observations suggest that polyprenol molecules prevent the formation of aggregated forms of LHCII, characterized by uncontrolled excitation quenching, although a certain ratio between polyprenols and other membrane components might be required to achieve this effect on protein aggregation. These results are in agreement with the observation done for isolated thylakoids that both deficiency and over-accumulation of polyprenols cause increased aggregation of LHCII (Fig. 5B).

### Polyprenols affect NPQ efficiency of the photosynthetic apparatus

Previously published studies on *cpt7* and *CPT7-OE* lines were referring only to standard growth conditions (Akhtar et al, 2017). Therefore, we decided to check if disturbances in polyprenol levels affect the responses of Arabidopsis plants to heat stress. Interestingly, we could not detect any changes in thermotolerance of neither *cpt7* nor *CPT7-OE* lines (Suppl. Fig. 7). Therefore, we investigated if polyprenols affect photosynthetic performance in response to high temperature. Various parameters reflecting photosystem I and photosystem II efficiency were measured *in vivo* using dark-adapted leaves. As previously reported (Akhtar et al, 2017), no significant differences in maximal quantum yield of PSII (F_v_/F_m_) were noticed (Suppl. Fig. 8A). The PSI quantum yield (Y(I)) measured in steady-state light conditions was significantly elevated in all heat-stressed plants with no visible differences between the genotypes (Suppl. Fig. 8B). However, we were unable to observe any substantial differences in PSII quantum yield across the tested genotypes and conditions, save for a slight increase for heat-treated *CPT7-OE* plants (Suppl. Fig. 8C). Moreover, no changes in the photochemical quenching parameter (qL), reflecting the rate of linear electron transport, could be observed between control WT, *cpt7* and *CPT7-OE* plants (Suppl. Fig. 8E), although heat stress caused a significant increase of qL in the *cpt7* mutant. Values of the non-photochemical quenching (NPQ) parameter obtained in steady-state light conditions did not differ significantly between control and heat-stressed plants across all genotypes (Suppl. Fig.8D). However, there was a clearly visible difference in the dynamics of NPQ curves registered in control or heat stress conditions for both WT and *CPT7-OE* plants, displaying much faster NPQ relaxation in high temperature (Fig. 6A and C). Interestingly, this difference disappeared for the *cpt7* line which displayed similar NPQ relaxation dynamics in both control and heat stress conditions (Fig. 6B). Furthermore, the relaxation pattern observed for *cpt7* was more akin to that observed for control WT plants than to the pattern for the heat-treated WT line (Fig. 6A).

**Figure 6.**
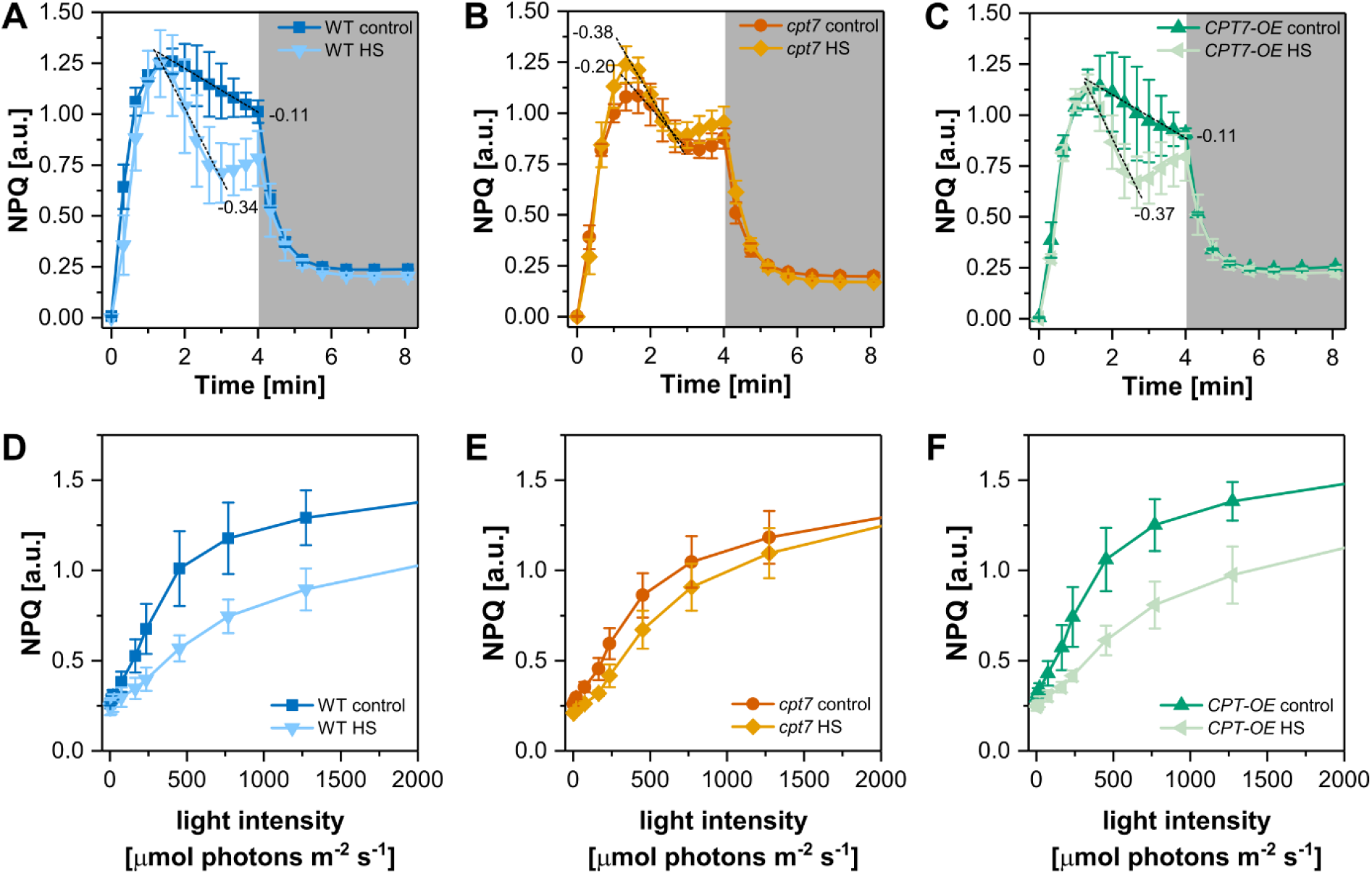
Polyprenols affect the function of Arabidopsis chloroplasts. **A-C** – changes in NPQ during the illumination of dark-adapted plants under high light conditions and during the dark recovery phase in WT (**A**), *cpt7* (**B**) and *CPT7-OE* (**C**) plants. Slopes of the observed curves are denoted. **D-F** – changes in NPQ under increasing light intensity in WT (**D**), *cpt7* (**E**) and *CPT7-OE* (**F**) plants. Leaves were light adapted (50 μmol photons s^−1^ m^−2^) and then subjected to illumination with increasing actinic light intensity (from 5 to 2000 μmol photons s^−1^ m^−2^). Means and SDs from three biological replicates are shown.

Analysis of the light curves recorded for light-adapted leaves revealed a significant difference in NPQ between *cpt7* and both WT and *CPT7-OE* plants. While values of NPQ were decreased after heat treatment in both WT and *CPT7-OE* plants (Fig. 6D and F), there was no significant difference between control and heat-stressed *cpt7* plants (Fig. 6E) and values in control conditions were lower than for the WT line. In case of the Y(I) and Y(II) responses, a slight increase after heat treatment was detected in all analyzed genotypes with no significant differences between them (Suppl. Fig. 9A-F). Values of qL were slightly increased after heat stress in WT and *cpt7* plants with no visible differences in the *CPT7-OE* line (Suppl. Fig. 9 G-I).

Taken together, the data presented above show that polyprenol deficiency affects the dynamics of NPQ changes during adaptation to light and modulates its response to heat treatment. These observations support the idea that polyprenols modulate chloroplast responses to external conditions.

## Discussion

### Plastidial polyprenols are environmentally regulated metabolites

Heat stress is one of the major environmental hazards affecting plants on many levels. It causes tissue dehydration, protein denaturation, disruption of lipid membranes and promotes the generation of reactive oxygen species. Lipids play a pivotal function in heat stress management. The key consequences of membrane lipid sensing of high temperature include membrane lipid rearrangement and generation of lipid signaling molecules as observed for yeast, plant and mammalian cells (Vigh et al, 1998; Mittler et al, 2012). Chloroplasts are especially susceptible to high temperature which affects photosynthesis at multiple levels causing chlorophyll degradation, PSII instability, disruption of electron transport and decreased Rubisco activity (Hu et al, 2020). To cope with the detrimental effects of heat stress, plants have evolved numerous mechanisms countering the effects of high temperature. These include the accumulation of chaperones, production of enzymes removing ROS and synthesis of ROS scavenging compounds like tocopherols and carotenoids (Kruk et al, 2003; Wang et al, 2018;).

Here, we show that synthesis of medium-chain length polyprenols (Pren-9, −10 and −11) that reside mainly in thylakoid membranes is considerably upregulated in response to heat stress. Accordingly, the expression of the gene encoding CPT7, the sole enzyme responsible for the synthesis of these compounds, is also induced in high temperature. Interestingly, there is a significant time gap between the peak of *CPT7* mRNA level and the onset of heat-induced production of polyprenols. Rapid return of the CPT7 transcript to the normal level suggests the involvement of posttranscriptional regulatory mechanisms that are yet to be identified. Moreover, upon recovery from heat stress, the polyprenol content decreases consistently. While this might be the result of polyprenol dilution in growing leaf tissue, it is also possible that a yet uncharacterized polyprenol catabolism pathway is involved.

While searching for regulators of *CPT7*, we have found that the induction of *CPT7* expression and accumulation of polyprenols depends on heat shock transcription factors belonging to the HSFA1 group. These master regulators of heat stress response in Arabidopsis were shown to govern the expression of numerous downstream-acting genes that are essential for adaptation to elevated temperature (Liu et al, 2011; Yoshida et al, 2011; Liu et al, 2013). In line with the presence of two non-canonical HSF binding sites (Guo et al, 2008) in the promoter sequence of *CPT7*, the results of ChIP lead us to consider *CPT7* a target of direct regulation by HSFA1s. According to our knowledge, HSFA1s are the first documented transcription factors that regulate expression of any *CPT* in Arabidopsis despite the fact that several reports show modulation of the expression of *CPTs* and polyisoprenoid biosynthesis in response to external cues. Studies on Arabidopsis hairy root cultures have shown that exogenously applied sugars and heavy metal ions as well as osmotic and oxidative stresses affect the expression of *CPTs* as well as the profile and content of dolichols (Jozwiak et al, 2013; Jozwiak et al, 2017). Moreover, the contribution of MVA- and MEP-derived IPP to dolichol biosynthesis changes upon osmotic stress (Jozwiak et al, 2017). Recently, it was shown that light enhances the production of polyprenols (Pren-9 to −12) in the leaves of *Tilia x euchlora* trees (Milewska-Hendel et al, 2017). Moreover, the content of long chain-polyprenols in leaves of trees and shrubs is affected by light conditions and changes throughout the vegetation season (Bajda et al, 2005). It should also be noted that medium-chain length polyprenols are present in multiple plant species tested, scattered across numerous botanical families (Swiezewska et al, 1994; Chouda and Jankowski, 2005). This shows that the presence of polyprenols is evolutionarily conserved and underlines their biological relevance.

### Polyprenols affect biochemical properties of chloroplasts

In the course of biochemical characterization of *cpt7* and *CPT7-OE* plants, we have found that disturbances in the polyprenol level affect, to some extent, the glycosylation status of chloroplast proteins. Our knowledge on the glycosylation of chloroplast proteins is very limited as only a handful of reports have been published on this topic (Asatsuma et al, 2005; Villarejo et al, 2005; Nanjo et al, 2006; Kitajima et al, 2009; Buren et al, 2011; Kaneko et al, 2016). It is known that chloroplast proteins undergo glycosylation in the ER/Golgi and then are transported to plastids (Villarejo et al,2005; Nanjo et al, 2006). The observed changes in glycosylation are most likely secondary effects resulting from altered chloroplast homeostasis. However, this observation requires further investigation.

Numerous reports indicate that plants adapt the lipid composition of their membranes to changing environmental conditions (Higashi and Saito, 2019; Shiva et al, 2020). The heat stress regime applied in our studies resulted in an increase in the levels of MGDG and DGDG. Although TEM analysis did not show any visible changes in the structural features of thylakoids that could accommodate the increased amount of membrane-forming lipids, we have observed an increased size of plastoglobules after heat treatment. Possibly, the increased pool of MGDG and DGDG observed in heat-treated plants is deposited in these structures. It should also be noted that the heat-induced increase in the level of polyprenols measured for isolated thylakoids was much lower than the increase observed for whole leaf tissue. This might suggest that not all polyprenols synthesized in response to heat stress are located in thylakoids. Possibly, a fraction of the synthesized polyprenols is deposited in some other plastidial compartment, e.g. the envelope, where their presence has been reported before (Akhtar et al, 2017).

### Polyprenols affect photosynthetic performance of chloroplasts

Interestingly, contrary to a previous report (Akhtar et al, 2017), our measurements of photosynthetic parameters in *cpt7* and *CPT7-OE* plants did not reveal significant changes in the steady state levels of PSII operating efficiency when compared to WT plants. Similarly, steady state photochemical quenching (qL) and non-photochemical quenching (NPQ) parameters remained unchanged while in the previous report polyprenol deficiency led to a decrease of qL and an increase of NPQ. These discrepancies might result from variations in plant growth conditions or from differences in the experimental methodology employed. This also implies that the biological functions of polyprenols are strongly influenced by environmental conditions. However, while in our hands the steady state level of the NPQ parameter remained unchanged across all tested genotypes, a time course measurement of NPQ revealed changes in its dynamics in the *cpt7* line. Similar results were obtained for light curves testing NPQ as a function of light intensity. Faster NPQ light-relaxation in *cpt7* control plants might be related to increased membrane fluidity of polyprenol-depleted thylakoid membranes as it might affect the diffusion rate of electron carriers (Akhtar et al, 2017).

In this study we have shown that the lack of polyprenols in the *cpt7* mutant causes a decrease in the stacking repeat distance (SRD) value. This could be the result of altered inter-complex interactions between adjacent thylakoids, causing tighter compaction of the stack and smaller partition gaps. However, the calculated parameter of stacking repeat distance does not precisely discriminate between changes in lumen volume of thylakoids and partition gap size, leaving open the possibility that the observed decrease in the SRD value results partly from lumen shrinking. We have also found that polyprenols affect the organization of protein complexes in thylakoid membranes. Studies on the fluorescence lifetime (FLIM) of chlorophyll molecules in LHCII complexes embedded in model membranes showed that polyprenols prevent protein aggregation. Accordingly, low-temperature (77 K) fluorescence measurements of thylakoid membranes isolated from *cpt7* plants revealed an increased content of aggregated LHCII. Interestingly, increased aggregation could also be observed in *CPT7-OE* plants, showing that a specific concentration of polyprenols in the lipid bilayer must be achieved to ensure proper aggregation levels. These observations support the notion that the lipid composition of thylakoid membranes affects the aggregation of LHCII, analogous to the disaggregating effect of PG and SQDG reported before (Schaller et al, 2011). Additionally, aggregated LHCII complexes can dissipate excitation energy in a manner similar to xanthophyll cycle-dependent NPQ (van Oort et al, 2007). Since polyprenols affect the aggregation of LHCII, they could possibly act as regulators of this form of energy dissipation. However, it should be noted that fluorescence spectra recorded for the *cpt7* and *CPT7-OE* lines indicated further changes in partitioning of LHCII complexes that differed between the analyzed genotypes, showing that absence or excess of polyprenols affect the organization of photosynthetic complexes in disparate ways.

Given that temperature conditions affect the partitioning of LHCs between different photosynthetic complexes (Nellaepalli et al, 2011; Nellaepalli et al, 2014), it might be possible that polyprenols promote or stabilize protein associations occurring during this process. It is postulated that polyisoprenoid alcohols can be bound by proteins with PIRS (polyisoprenoid recognition sequence) domains such as in eukaryotic and bacterial glycosyltransferases (Albright et al, 1989; Hartley and Imperial, 2012). A single lipid can be bound by multiple PIRS peptides which could facilitate the formation of multiprotein complexes (Hartley and Imperiali, 2012). Analysis of lipid-protein interactions between polyprenols and components of the photosynthetic apparatus could shed some light on this subject. Somewhat surprisingly, the effect of polyprenols on membrane fluidity and LHCII aggregation was considerably weaker in heat-stressed plants than in plants grown in standard conditions. This indicates that multiple stress-responsive mechanisms may operate in parallel in Arabidopsis cells to secure appropriate adaptation to heat. In our studies polyprenol levels remained strongly elevated several days after heat treatment, indicating a possible role for polyprenols in maintaining heat stress memory. This intriguing possibility deserves further studies.

## Conclusions

The role of plastidial polyprenols is an emerging field of study requiring more in-depth research on the mechanical aspects of their mode of action. While most chloroplast studies ignore polyprenols, we present evidence that these compounds are one of the major constituents of thylakoid membranes. This and previous reports (Bykowski et al, 2020; Van Gelder et al, 2018; Akhtar et al, 2017) have shown that polyprenols subtly but clearly affect many aspects of chloroplast biology. We believe that these compounds act as fine-tuning agents that help plants in adapting to environmental conditions. The biosynthesis of polyisoprenoids responds to external cues and might be one of the many links connecting the perception of heat to adaptive mechanisms that allow plants to withstand unfavorable growth conditions. Although further studies on these not well-known compounds are indeed required, the results obtained so far give premise for better understanding of plastidial biology and, possibly, for novel biotechnological approaches to obtaining plant varieties that could cope with ongoing climate changes.

## Acknowledgements

Authors are grateful to Dr Marta Hoffman-Sommer for help with manuscript preparation and editing. This work was supported by National Science Centre, Poland grants UMO-2017/26/D/NZ1/00833 (to D.B.), UMO-2018/29/B/NZ3/01033 (to E.S.) and UMO-2016/21/B/NZ1/02793 (to L.S.). Authors declare no conflicts of interest.

